# Genetic evidence for shared risks across psychiatric disorders and related traits in a Swedish population twin sample

**DOI:** 10.1101/234963

**Authors:** Mark J. Taylor, Joanna Martin, Yi Lu, Isabell Brikell, Sebastian Lundström, Henrik Larsson, Paul Lichtenstein

**Author notes:** joint first authors (Mark J. Taylor, Joanna Martin).

## Abstract

Psychiatric traits related to categorically-defined psychiatric disorders are heritable and present to varying degrees in the general population. In this study, we test the hypothesis that genetic risk factors associated with psychiatric disorders are also associated with continuous variation in milder population traits. We combine a contemporary twin analytic approach with polygenic risk score (PRS) analyses in a large population-based twin sample. Questionnaires assessing traits of autism spectrum disorder (ASD), attention-deficit/hyperactivity disorder (ADHD), learning difficulties, tic disorders (TD), obsessive-compulsive disorder (OCD), anxiety, major depressive disorder (MDD), mania and psychotic experiences were administered to a large, Swedish twin sample. Individuals with clinical psychiatric diagnoses were identified using the Swedish National Patient Register. Joint categorical/continuous twin modeling was used to estimate genetic correlations between psychiatric diagnoses and continuous traits. PRS for psychiatric disorders were calculated based on independent discovery genetic data. The association between PRS for each disorder and related continuous traits was tested. We found mild to strong genetic correlations between psychiatric diagnoses and corresponding traits (ranging from .31-.69) in the twin analyses. There was also evidence of association between PRS for ASD, ADHD, TD, OCD, anxiety, MDD and schizophrenia with related population traits. These results indicate that genetic factors which predispose to psychiatric disorders are also associated with milder variation in characteristic traits throughout the general population, for many psychiatric phenotypes. This finding supports the conceptualization of psychiatric disorders as the extreme ends of continuous traits.

## Introduction

Psychiatric disorders are highly impairing and strongly associated with genetic factors^1^. They are diagnosed on the basis of an individual meeting a number of specific symptoms that are associated with other clinical features (e.g. duration, distress, impairment, and onset). Individuals not meeting diagnostic criteria for a particular disorder can nonetheless present with behaviors characteristic of that disorder. Indeed, continuously-distributed traits relevant to psychiatric symptoms are present to varying degrees throughout the general population and are often as heritable as the clinical disorder^2^. One hypothesis is that psychiatric disorders share genetic risks with such continuously-distributed traits in the general population^2^. A few studies using classical twin and more recent molecular genetic methods have lent preliminary support to this hypothesis for several psychiatric phenotypes.

Twin studies have used DeFries-Fulker analysis to estimate group heritability; significant group heritability suggests genetic continuity between milder psychiatric traits and more severe manifestations^3^. Significant group heritability has been reported for extreme traits of autism spectrum disorder (ASD), attention-deficit/hyperactivity disorder (ADHD), learning difficulties, and anxiety^4^. While DeFries-Fulker analysis can test for quantitative similarities in the etiology of mild and severe traits, it does not yield a genetic correlation between them. A more contemporary approach was taken in a UK-based twin study, which reported a genetic correlation of .70 between best-estimate diagnoses of ASD and autistic traits^5^. This approach has not been applied to other psychiatric phenotypes.

Genetic correlation estimates based on genome-wide common variants support a high degree of shared risks across disorders and traits for ADHD and MDD, with a more moderate estimate for ASD^4^. Robust estimates of common variant genetic correlations using available methods^6,7^ require very large genome-wide datasets, which are not yet available for many psychiatric phenotypes. An alternative method is to leverage discovery genome-wide association studies (GWAS) of psychiatric disorders for polygenic risk score (PRS) analysis^8^. Preliminary studies have reported associations between psychiatric disorder PRS with related population traits for ASD, ADHD, obsessive-compulsive disorder (OCD), and major depressive disorder (MDD), with null or mixed findings for schizophrenia and bipolar disorder^4^.

Although there is preliminary evidence for shared genetic risks across disorder and traits for some phenotypes, the evidence is either weak, mixed or entirely lacking for many psychiatric phenotypes. Consequently, we aimed to assess the degree to which the genetic factors associated with clinical diagnoses of psychiatric disorders are also associated with continuous variation in traits of the same phenotype, for a range of psychiatric disorders. We used data from a large, population-based twin sample, which has been genotyped and linked with the Swedish National Patient Register^9^. This unique data source allowed us to combine twin modelling with molecular genetic methods in the same sample. Specifically, we first estimated genetic correlations between psychiatric disorders and continuous traits using twin methods. We then tested whether PRS for psychiatric disorders derived from recent GWAS are associated with continuous variation in related population traits.

## Method

### Participants

Families of all twins born in Sweden since 1992 are contacted in connection with the twins’ 9^th^ birthday (the first three years of the study also included 12 year olds), and invited to participate in the ongoing Child and Adolescent Twin Study in Sweden (CATSS)^10^. The response rate at first contact is 75%. Families are then followed up when the twins are aged 15 (61% response rate) and i8 (59% response rate). Exclusion criteria for the current study were brain injuries (N=207 pairs), chromosomal syndromes (N=35 pairs), death (N=29 pairs), and migration (N=100 pairs). Phenotypic data were available for 13,923 pairs at age 9 or 12, 5,165 pairs at age 15, and 4,273 pairs at age 18. Zygosity was ascertained using a panel of 48 single nucleotide polymorphisms (SNPs) or an algorithm of five questions concerning twin similarity. Only twins with a 95% probability of correct classification have been assigned zygosity using the latter method. Zygosity was reconfirmed for pairs with genotype data available. This study has ethical approval from the Karolinska Institutet Ethical Review Board.

DNA samples (from saliva) were obtained from the CATSS participants at study enrollment. N=11,551 individuals with available DNA were genotyped using the Illumina PsychChip. Standard quality control (QC) and imputation procedures were performed in the sample; for details see Brikell et al.^11^. N=11,081 samples passed QC; MZ twins were then imputed resulting in N=13,576 samples and N=6,981,993 imputed SNPs passing all QC. After individual-level exclusions (described above), N=13,412 children were included in genetic analyses.

### Phenotypic measures

#### Clinical Diagnoses

CATSS has been linked with the Swedish National Patient Register (NPR)^9^. The NPR contains records of specialist in-patient and out-patient care in Sweden, and includes ICD-10 diagnoses for each visit to such services. For CATSS, in-patient data were available between 1987-2014, and out-patient data between 2001-2013. Diagnoses of ASD, ADHD, intellectual disability (ID), tic disorders (TD), OCD, anxiety disorders (ANX), and MDD were extracted. All diagnoses were treated dichotomously. Diagnostic codes and the numbers of individuals with each diagnosis for twin and genetic analyses are shown in Table 1.

**Table 1:** Description of the CATSS sample and measures

| Phenotype | ICD Diagnostic Codes | NPR Diagnosis |  | Study Measures | Description of Measure | N of individuals with data (twin analyses) | N of individuals with data (genetic analyses) |
| --- | --- | --- | --- | --- | --- | --- | --- |
|  |  | N (%) Affected (twin analyses) | N (%) Affected (genetic analyses) |  |  |  |  |
| **ASD** | F84 | 253 (0.91%) | 142 (1.06%) |  | <u>Age 9/12</u>: Parent-rated A-TAC ASD module (17 items) | 27,780 | 13,396 |
| **ADHD** | F90 | 824 (2.96%) | 440 (3.28%) |  | <u>Age 9/12</u>: Parent-rated A-TAC ADHD module (19 items) | 27,759 | 13,391 |
| **ID** | F70-F73 | 166 (0.60%) | 77 (0.57%) |  | <u>Age 9/12</u>: Parent-rated A-TAC Learning module (3 items) | 27,804 | 13,400 |
| **TD** | F95 | 103 (0.37%) | 43 (0.32%) |  | <u>Age 9/12</u>: Parent-rated A-TAC Tics module (3 items) | 27,791 | 13,396 |
| **OCD** | F42 | 118 (0.42%) | 67 (0.50%) |  | <u>Age 9/12</u>: Parent-rated ATAC Compulsions module (2 items);<br><u>Age 18</u>: Self-rated BOCS (15 items) | 27,802;<br>5,757 | 13,400;<br>3,982 |
| **ANX** | F40-F41 | 474 (1.70%) | 251 (1.87%) |  | <u>Age 9/12</u>: Parent-rated SCARED (41 items);<br><u>Age 15</u>: Parent- and self-rated SDQ Emotion subscale (5 items);<br><u>Age 18</u>: Parent-rated ABCL DSM-IV Anxiety subscale (6 items);<br>Self-rated SCARED (38 items) | 15,589;<br>7,663;<br>8,150;<br>5,160;<br>5,801 | 6,806;<br>5,703;<br>5,917;<br>3,755;<br>4,007 |
| **MDD** | F32-F34 | 414 (1.49%) | 222 (1.66%) |  | <u>Age 9/12</u>: Parent-rated SMFQ (13 items);<br><u>Age 15</u>: Parent- and self-rated SDQ Emotion subscale (5 items);<br><u>Age 18</u>: Parent-rated ABCL DSM-IV Depression subscale (15 items);<br>Self-rated CES-D (11 items) | 15,873;<br>7,663;<br>8,150;<br>5,215;<br>5,518 | 6,826;<br>5,703;<br>5,917;<br>3,791;<br>3,813 |
| **Mania** | N/A | N/A | N/A |  | <u>Age 18</u>: Parent- and self-rated MDQ (13 items) | N/A | 3,808;<br>4,128 |
| **Psychosis** | N/A | N/A | N/A |  | <u>Age 18</u>: Parent- and self-rated APSS (7 items) | N/A | 5,368;<br>5,518 |

ASD: autism spectrum disorders; ADHD: attention-deficit/hyperactivity disorder; ID: intellectual disability; TD: tic disorders; OCD: obsessive-compulsive disorder; ANX: anxiety disorders; MDD: major depressive disorder; ABCL: Adult Behavior Checklist<sup>12</sup>; A-TAC: Autism-Tics, AD/HD, and other Comorbidities inventory<sup>13,14</sup>; APSS: Adolescent Psychotic-Like Symptoms Screener<sup>15</sup>; BOCS: Brief Obsessions and Compulsions Scale<sup>16</sup>; CES-D: Center for Epidemiologic Studies Depression scale<sup>17</sup>; MDQ: Mood Disorders Questionnaire<sup>18</sup>; NPR: National Patient Register; SCARED: Screen for Anxiety Related Disorders<sup>19</sup>; SDQ: Strengths and Difficulties Questionnaire<sup>20</sup>; SMFQ: Short Moods and Feelings Questionnaire<sup>21</sup>.
N.B. Diagnoses of bipolar disorder and schizophrenia were not included due to small numbers of participants with these diagnoses. There were too few items to divide the SDQ Emotion subscale into anxiety and depression separately.

#### Continuous Measures

Traits of ASD, ADHD, ID, TD, OCD, ANX and MDD were measured using continuous scales at ages 9/12 years. Internalizing problems (related to ANX and MDD) were then measured at age 15. Traits of OCD, ANX, MDD, mania, and psychotic-like experiences were assessed when the twins were followed up at age 18. Details of these measures, along with sample sizes, are provided in Table 1.

### Analyses

#### Twin Analyses

We used joint categorical-continuous trait twin models to assess the degree of etiological overlap between continuous traits and categorical diagnoses. For each phenotype, the model estimates the degree of variation attributable to additive genetic (A), non-additive genetic (D), shared environmental (C), and non-shared environmental influences (E; with the non-shared environmental component encompassing measurement error). It then estimates the correlations between these components across the two modeled phenotypes. The phenotypic correlation is then decomposed into genetic and environmental influences, thus assessing which etiological factors lead to continuous traits being correlated with categorical psychiatric disorders. Based on twin correlations, we tested either ACE or ADE models for each disorder-trait pairing. Where an ADE model was tested, we also modeled sibling interaction effects (termed ‘s’), since these effects can mimic the effects of D on the twin correlations^22^. The general principles of the twin design are described at length elsewhere^23^.

ACE or ADE-s models were fitted based on the twin correlations, and compared with a baseline saturated model of the observed data. Where twin correlations were inconsistent (i.e. the pattern of correlations differed between the continuous trait and the categorical diagnosis), both ACE and ADE-s models were fitted and the best-fitting model was selected on the basis of the lowest Bayesian Information Criteria (BIC) value. In order to test more parsimonious explanations of the data, each model was reduced by constraining certain components to equal zero. These nested models were compared to the full ACE or ADE-s model using the likelihood-ratio test; if no significant deterioration of model fit was observed, then the reduced model was favored. All continuous scales were standardized by sex, while the effects of sex on the thresholds were included in the models. All models were fitted in OpenMx^24^. We included opposite-sex twins but did not test for quantitative sex differences due to insufficient power, and assumed no qualitative sex differences. OCD was omitted from the twin analyses due to the small sample and low heritability.

#### Polygenic risk score analyses

Publicly available GWAS summary statistics for 8 psychiatric disorders (i.e. ASD, ADHD, TD, OCD, ANX, MDD, bipolar disorder (BD) and schizophrenia) and 3 continuously distributed psychiatric or cognitive traits (i.e. ADHD symptoms, cognitive ability and depressive symptoms) were used to derive PRS in the CATSS individuals^25–34^. Table S1 lists each of these discovery datasets along with sample sizes. See Supplemental Text for details of how PRS were derived. In brief, PRS were calculated for each individual by scoring the number of effect alleles (weighted by the SNP effect size) across each discovery set of clumped SNPs in PLINK.v.1.9, for a range of p-value thresholds used for SNP selection. The primary analyses are based on the threshold p<0.5. The PRS were standardized using z-score transformations; effect sizes can be interpreted as increase in risk of the outcome, per standard deviation increase in PRS. Principal component analysis was used to derive covariates to account for population stratification (Supplemental Text).

Analyses of PRS in relation to outcomes were performed using generalized estimating equations (GEE) implemented in the R package ‘drgee’, with robust standard errors, based on clustering related individuals according to family IDs. Principal components, sex and age (for measures that were assessed at age 9 or 12 years) were included as covariates. First, we tested for association between PRS for each of the eight discovery GWAS of psychiatric disorders and the corresponding continuously distributed population trait(s). Second, these analyses were repeated after excluding individuals diagnosed with the relevant psychiatric disorder, based on available information on ICD diagnoses. This was done to determine whether effects were driven primarily by individuals with clinically-recognized problems. Third, we tested for associations between PRS for each of the three discovery GWAS of continuously-distributed population traits and presence of the corresponding psychiatric diagnosis in the target sample. Finally, all PRS analyses were repeated using PRS derived based on different p-value selection thresholds, to assess sensitivity. False discovery rate (FDR) corrections were applied in R (using the “fdr” method, in the function “p.adjust”) to account for multiple testing.

#### Supplemental Analyses

Since there is some evidence of genetic specificity within ASD and ADHD trait domains^35,36^, we also re-ran all twin and PRS analyses for three specific DSM-IV ASD dimensions (social problems, language impairment, and behavioral inflexibility) and two DSM-IV ADHD dimensions (hyperactivity/impulsivity and inattention). These domains were assessed by dividing the A-TAC ASD and ADHD subscales, based on prior publications^10^.

## Results

### Twin Analyses

The probandwise concordances for each ICD diagnosis, which represent the probability of the cotwin of a proband also receiving a given diagnosis, are presented in Table S2. The MZ probandwise concordances exceeded the DZ probandwise concordances, indicating genetic influences on each diagnosis. The phenotypic and twin correlations are shown in Table 2. Phenotypic correlations between disorders and related traits ranged from modest (.22 for MDD) to strong (.66 for ID). MZ twin correlations were higher than DZ correlations within each disorder and trait, and across disorder-trait phenotype pairs, indicating that each clinical diagnosis and continuous trait, as well as the covariance between them, was influenced by genetic factors.

**Table 2:**
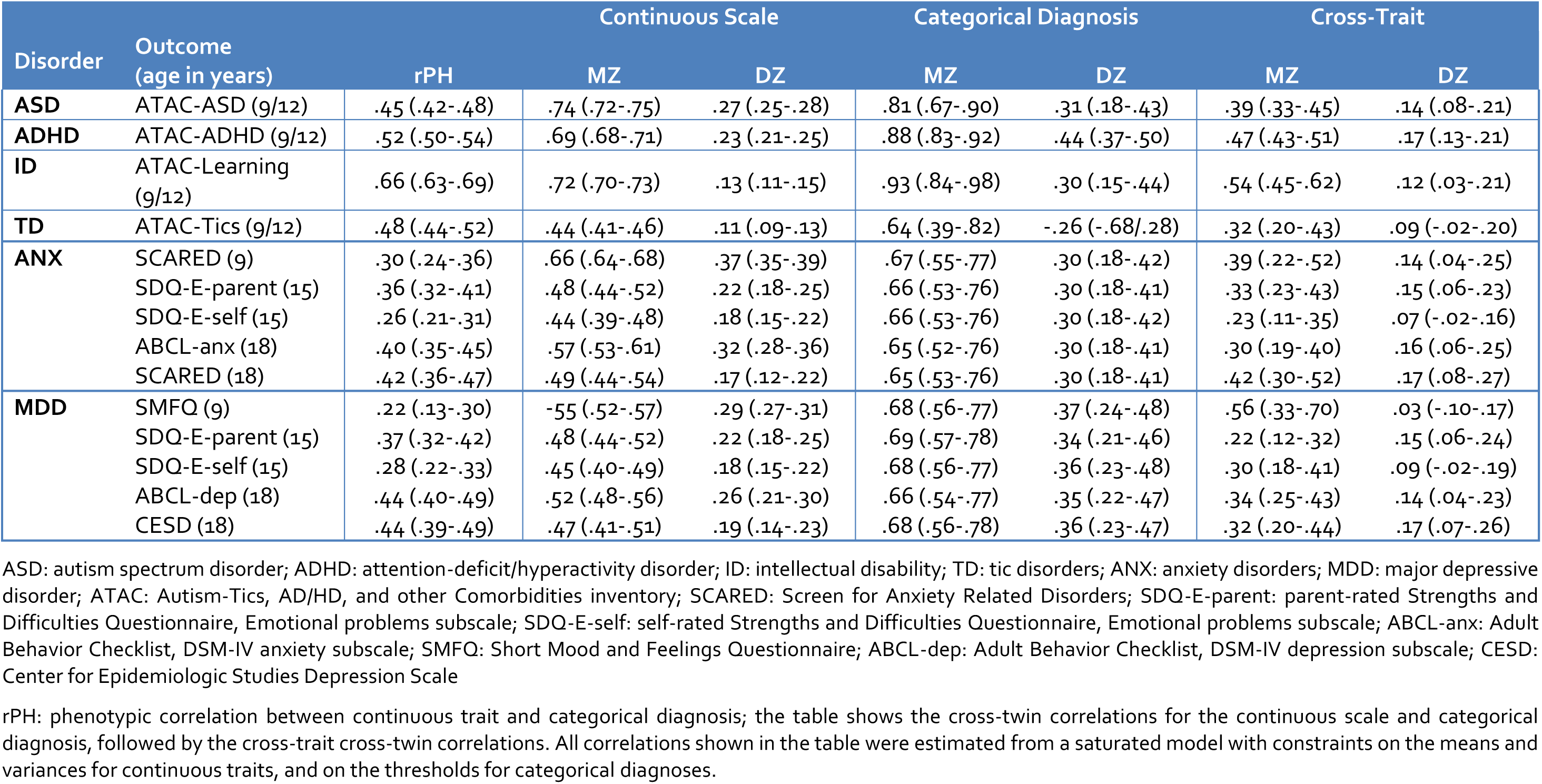
Phenotypic and twin correlations across disorders and continuous traits

The joint categorical-continuous twin model fit statistics are shown in Table S3. Where ADE-s models were fitted, the D component could be dropped. For traits with ACE models, shared environment was non-significant for all traits, except for anxiety at age 9/12, where C played a modest role.

Parameter estimates from the best-fitting model for each phenotype are shown in Table S4. For all measures and diagnoses, except for the SDQ at age 15 and self-reported MDD at age 18, genetic factors were more influential than environmental factors. The genetic and non-shared environmental correlations are shown in Figure 1. Estimates of genetic correlation varied in magnitude: .48 for ASD, .56 for ADHD, .69 for ID, .61 for TD, .46-.57 for anxiety, and .33-.58 for MDD. For anxiety and MDD, higher genetic correlations were observed at age 18. These shared genetic influences were the key factor explaining the correlations between traits and diagnoses, as shown in Table S4; 60-100% of the correlation between each trait and diagnosis was explained by additive genetics.

**Figure 1:**
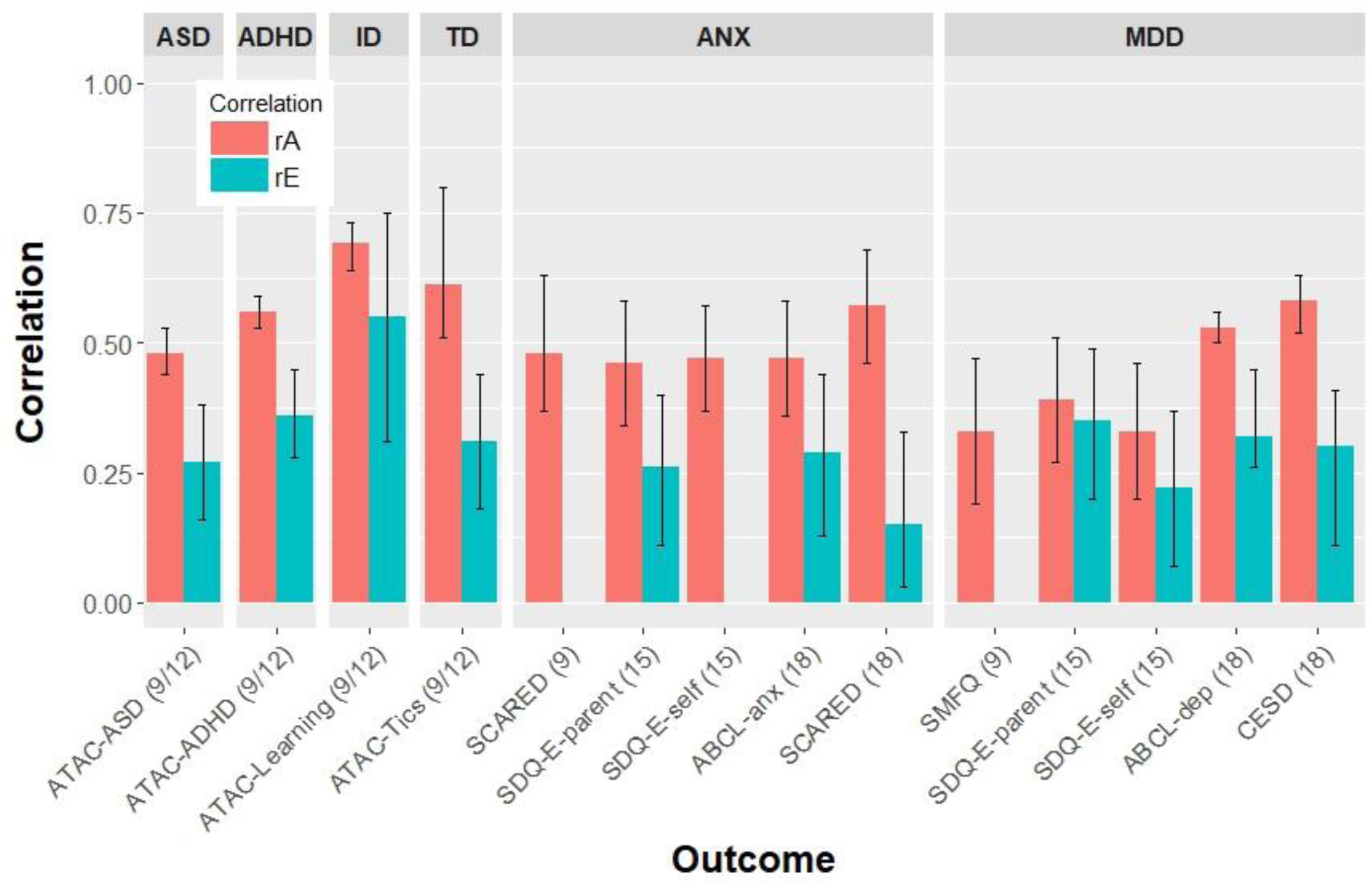
Genetic and nonshared environmental correlations between psychiatric diagnoses and continuous traits from the best-fitting twin models. ASD: autism spectrum disorder; ADHD: attention-deficit/hyperactivity disorder; ID: intellectual disability; ANX: anxiety disorders; MDD: major depressive disorder A-TAC: Autism-Tics, AD/HD, and other Comorbidities inventory; SCARED: Screen for Anxiety Related Disorders; SDQ-E-parent: parent-rated Strengths and Difficulties Questionnaire, Emotional problems subscale; SDQ-E-self: self-rated Strengths and Difficulties Questionnaire, Emotional problems subscale; ABCL-anx: Adult Behavior Checklist, DSM-IV anxiety subscale; SMFQ: Short Mood and Feelings Questionnaire; ABCL-dep: Adult Behavior Checklist, DSM-IV depression subscale; CESD: Center for Epidemiologic Studies Depression Scale. rA: genetic correlation; rE: non-shared environmental correlation

The additional analyses of specific ASD and ADHD domains are shown in Tables S5-S6. The results were consistent across trait dimensions. All three ASD trait domains displayed moderate phenotypic (.43-.48) and genetic correlations (.43-.47) with ASD, while both ADHD dimensions correlated moderately with ADHD, both phenotypically (.45-.53) and genetically (.49-.57).

### PRS analyses

PRS for clinically-defined psychiatric disorders were associated with related quantitative traits, for the following phenotypes: ASD, ADHD, TD, OCD, ANX, MDD and schizophrenia (Table 3). No association was seen for BD. OCD PRS were associated with obsessive-compulsive symptoms at age 18 but not at age 9/12. PRS for ANX were associated with anxiety traits across different time points (ages 9-18 years), with the exception of self-rated internalizing traits at age 15 and parent-rated symptoms at age 18. MDD PRS were associated with depressive symptoms at ages 15 and 18 but not at age 9. After removing individuals diagnosed with the relevant psychiatric disorder based on NPR diagnoses, the results remained significant for ASD, ADHD, TD, OCD, anxiety and MDD though the effect size decreased and was no longer significant for anxiety at age 18 (Table 3).

**Table 3:**
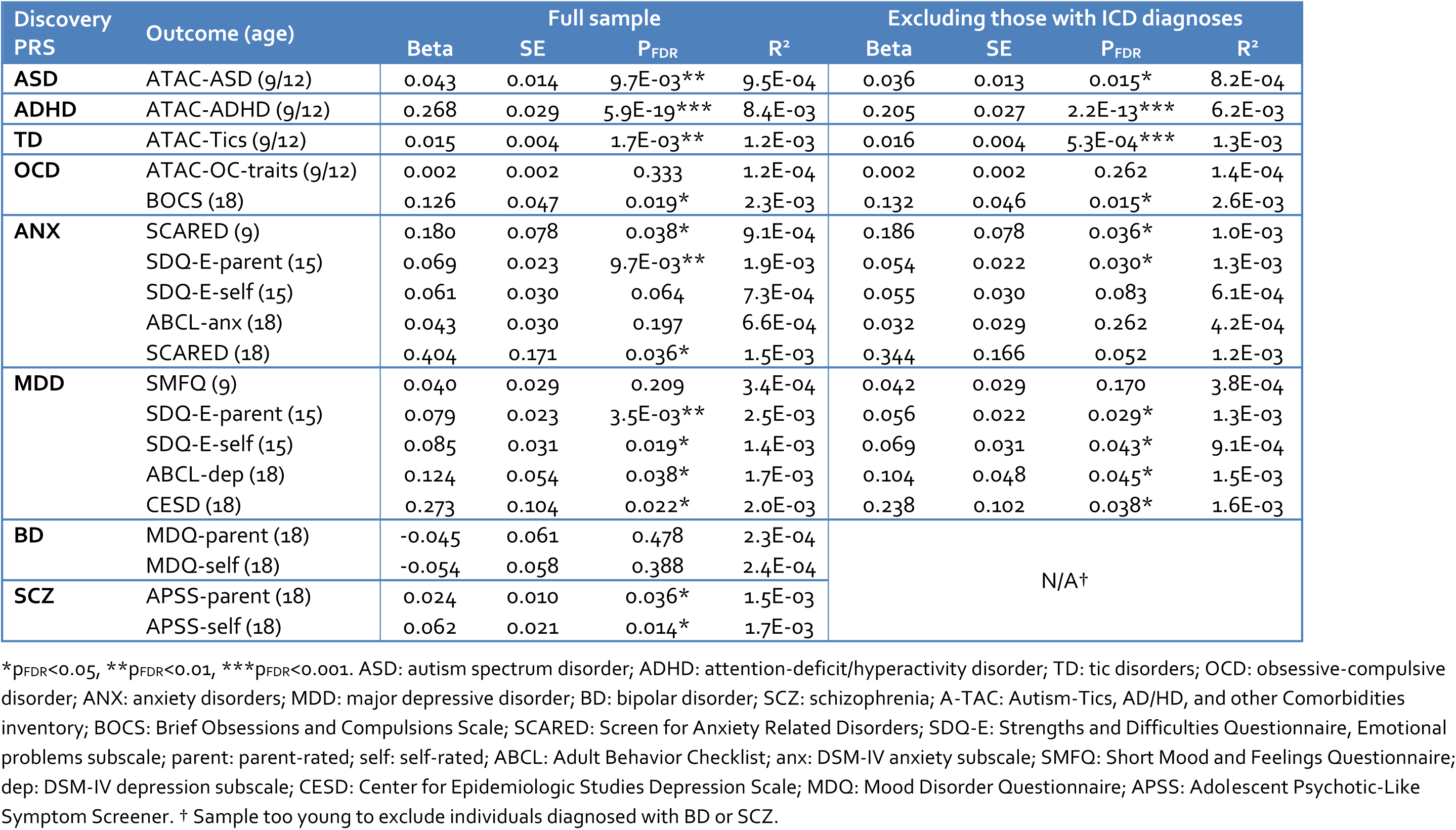
Association of polygenic risk scores with related continuous outcomes

| Discovery PRS | Outcome (age) | Full sample |  |  |  | Excluding those with ICD diagnoses |  |  |  |
| --- | --- | --- | --- | --- | --- | --- | --- | --- | --- |
|  |  | Beta | SE | P<sub>FDR</sub> | R<sup>2</sup> | Beta | SE | P<sub>FDR</sub> | R<sup>2</sup> |
| **ASD** | ATAC-ASD (9/12) | 0.043 | 0.014 | 9.7E-03** | 9.5E-04 | 0.036 | 0.013 | 0.015* | 8.2E-04 |
| **ADHD** | ATAC-ADHD (9/12) | 0.268 | 0.029 | 5.9E-19*** | 8.4E-03 | 0.205 | 0.027 | 2.2E-13*** | 6.2E-03 |
| **TD** | ATAC-Tics (9/12) | 0.015 | 0.004 | 1.7E-03** | 1.2E-03 | 0.016 | 0.004 | 5.3E-04*** | 1.3E-03 |
| **OCD** | ATAC-OC-traits (9/12) | 0.002 | 0.002 | 0.333 | 1.2E-04 | 0.002 | 0.002 | 0.262 | 1.4E-04 |
|  | BOCS (18) | 0.126 | 0.047 | 0.019* | 2.3E-03 | 0.132 | 0.046 | 0.015* | 2.6E-03 |
| **ANX** | SCARED (9) | 0.180 | 0.078 | 0.038* | 9.1E-04 | 0.186 | 0.078 | 0.036* | 1.0E-03 |
|  | SDQ-E-parent (15) | 0.069 | 0.023 | 9.7E-03** | 1.9E-03 | 0.054 | 0.022 | 0.030* | 1.3E-03 |
|  | SDQ-E-self (15) | 0.061 | 0.030 | 0.064 | 7.3E-04 | 0.055 | 0.030 | 0.083 | 6.1E-04 |
|  | ABCL-anx (18) | 0.043 | 0.030 | 0.197 | 6.6E-04 | 0.032 | 0.029 | 0.262 | 4.2E-04 |
|  | SCARED (18) | 0.404 | 0.171 | 0.036* | 1.5E-03 | 0.344 | 0.166 | 0.052 | 1.2E-03 |
| **MDD** | SMFQ (9) | 0.040 | 0.029 | 0.209 | 3.4E-04 | 0.042 | 0.029 | 0.170 | 3.8E-04 |
|  | SDQ-E-parent (15) | 0.079 | 0.023 | 3.5E-03** | 2.5E-03 | 0.056 | 0.022 | 0.029* | 1.3E-03 |
|  | SDQ-E-self (15) | 0.085 | 0.031 | 0.019* | 1.4E-03 | 0.069 | 0.031 | 0.043* | 9.1E-04 |
|  | ABCL-dep (18) | 0.124 | 0.054 | 0.038* | 1.7E-03 | 0.104 | 0.048 | 0.045* | 1.5E-03 |
|  | CESD (18) | 0.273 | 0.104 | 0.022* | 2.0E-03 | 0.238 | 0.102 | 0.038* | 1.6E-03 |
| **BD** | MDQ-parent (18) | -0.045 | 0.061 | 0.478 | 2.3E-04 | N/A† |  |  |  |
|  | MDQ-self (18) | -0.054 | 0.058 | 0.388 | 2.4E-04 |  |  |  |  |
| **SCZ** | APSS-parent (18) | 0.024 | 0.010 | 0.036* | 1.5E-03 |  |  |  |  |
|  | APSS-self (18) | 0.062 | 0.021 | 0.014* | 1.7E-03 |  |  |  |  |

\*p<sub>FDR</sub><0.05, \*\*p<sub>FDR</sub><0.01, \*\*\*p<sub>FDR</sub><0.001. ASD: autism spectrum disorder; ADHD: attention-deficit/hyperactivity disorder; TD: tic disorders; OCD: obsessive-compulsive disorder; ANX: anxiety disorders; MDD: major depressive disorder; BD: bipolar disorder; SCZ: schizophrenia; A-TAC: Autism-Tics, AD/HD, and other Comorbidities inventory; BOCS: Brief Obsessions and Compulsions Scale; SCARED: Screen for Anxiety Related Disorders; SDQ-E: Strengths and Difficulties Questionnaire, Emotional problems subscale; parent: parent-rated; self: self-rated; ABCL: Adult Behavior Checklist; anx: DSM-IV anxiety subscale; SMFQ: Short Mood and Feelings Questionnaire; dep: DSM-IV depression subscale; CESD: Center for Epidemiologic Studies Depression Scale; MDQ: Mood Disorder Questionnaire; APSS: Adolescent Psychotic-Like Symptom Screener. † Sample too young to exclude individuals diagnosed with BD or SCZ.

In the analysis of PRS for quantitative traits in relation to psychiatric diagnoses (Table 4), PRS for depressive symptoms were associated with MDD diagnosis; PRS for traits of ADHD and general cognitive ability were not associated with related diagnoses (ADHD and ID, respectively).

**Table 4:**
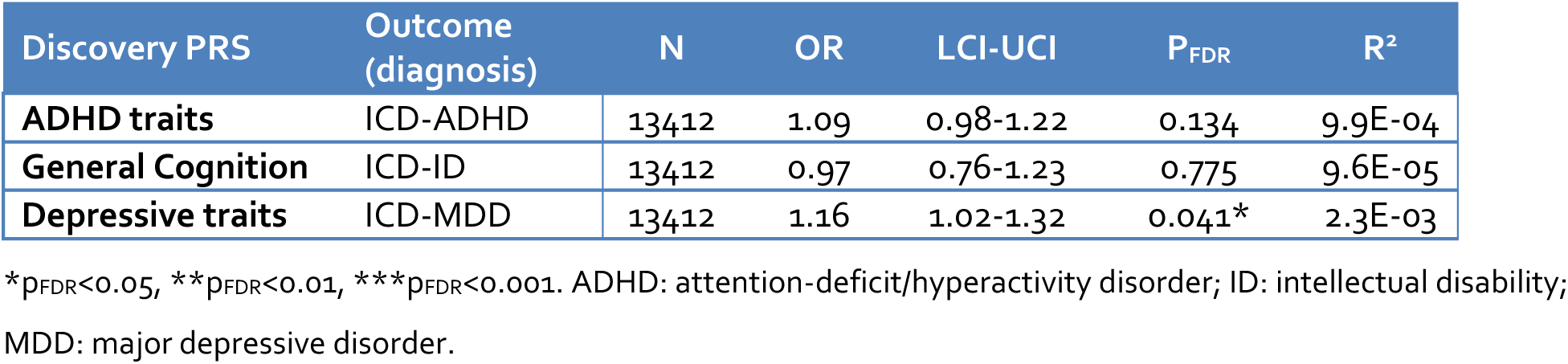
Association of continuous trait polygenic risk scores with related binary outcomes

Secondary analyses for ASD and ADHD PRS in relation to traits of specific domains are shown in Table S7. The results of analyses of these sub-domains were consistent with the analyses of total ASD and ADHD trait scores, except that for ASD, only the estimate for the domain of flexibility remained significant after the exclusion of individuals with ASD diagnoses.

All main analyses were repeated using PRS derived based on different p-value thresholds for SNP inclusion (Figure S1). The pattern of results was reasonably consistent in these sensitivity analyses, albeit due to the additional tests performed, several results (e.g. for anxiety at ages 9 and 18) were not statistically significant after FDR correction.

## Discussion

In this paper, we leveraged data from a large twin sample with information on psychiatric diagnoses and traits as well as common variant genetic data, to test the degree of genetic association between clinical psychiatric diagnoses and related traits. For all disorders analyzed (i.e. ASD, ADHD, TD, ID, ANX and MDD) the twin modelling results indicated significant, and sometimes strong, genetic correlations between the clinical diagnosis and related trait(s). These results were supported by common variant PRS analyses, with robust evidence of shared risks between disorders and traits seen for ASD, ADHD, TD, OCD, ANX, MDD and schizophrenia, even after exclusion of diagnosed individuals, where possible. Furthermore, PRS analyses showed no clear evidence of associations for BD or ID. These results provide the most comprehensive evidence to date that many psychiatric disorders can be considered as extreme manifestations of one or more continuous symptom dimensions in the population.

Our study extends the existing literature by directly estimating the genetic correlation between psychiatric disorders and continuous traits based on twin data, while also assessing the association of genetic risk across disorders and traits using PRS analysis in the same sample. It is becoming increasingly clear that considering psychiatric disorders as dichotomous may not be the optimal approach for all research studies of these phenotypes. In terms of molecular genetic research, these results support the need for novel methods that jointly analyze psychiatric disorders and continuously measured psychiatric traits in order to improve statistical power and yield insights into the biology of these phenotypes. Indeed, the value of such an approach was recently demonstrated in an ADHD GWAS, which yielded additional genome-wide significant loci when meta-analyzing case-control and trait GWAS data^26^. Genetic studies of continuous traits in community-based samples may also have the added benefit of being more representative than clinical samples, while also generating results that could be generalized to clinical populations.

It is important to note that the genetic correlations in the twin analyses were estimated to be less than 1, and associations between PRS and traits had modest effect sizes. These results could be related to the relatively young age of the sample. For example, the genetic correlation between MDD and depressive traits was .33 at ages 9 and 12, but increased to .58 at age 18. Another possible interpretation of these results is that a proportion of genetic influences on any given psychiatric disorder is unique and not shared with milder related population traits. This may be especially true for rare genetic variants which are likely to have a more deleterious phenotypic impact; however, several studies have demonstrated some degree of genetic overlap from rare variants across disorders and continuous measures of ASD^37^ and ID^38^. Future work identifying genetic risk variants that are not shared between psychiatric traits and more severe clinical manifestations may help elucidate why some individuals present only with mild traits, while others go on to develop clinically severe problems. Related to this point, it is important to stress that this study did not examine several important clinical features associated with psychiatric disorders, such as symptom duration, timing, persistence, impairment, or distress. Therefore, the results of this study do not imply that mild psychiatric traits are necessarily harmful.

Our results should be interpreted in light of several strengths as well as a number of caveats. It is a major strength that we were able to perform both twin and molecular genetic analyses in a single cohort, owing to our unique sample of twins with both diagnostic and genetic data. This unique data source also allowed for the exclusion of individuals diagnosed with psychiatric disorders from the PRS analyses, to allow for a more robust test ruling out the possibility that observed effects were only driven by individuals with clinically-recognized problems. This sample has been assessed at multiple ages, using both parental- and self-reports, leading to a wealth of information on psychiatric phenotypes. Nonetheless, the sample is still rather young. While many disorders have typically developed during childhood and by emerging adulthood, other disorders are more common at later ages. We therefore could not perform twin analyses of the associations between schizophrenia or BD and related traits, or exclude individuals with these diagnoses from the PRS analyses, since very few of our participants had passed through the risk age for these disorders. Similarly, analyses of individuals diagnosed with MDD may have missed individuals who will go on to be diagnosed with MDD in the future. It is also important to note that the National Patient Register covers only specialist care, and so individuals with psychiatric disorders who were treated only in primary care settings were likely missed.

Furthermore, we did not have the statistical power to divide our clinical cases by severity or diagnostic subtype. Thus, we cannot conclude that all levels of disorder severity share genetic risks with milder traits. This will be an important focus in future research, since there is some evidence that very severe ID is genetically independent from cognitive abilities and milder ID^39^. Also, all the continuous measures used in this study were designed to assess potentially problematic behaviors. As such, we cannot extrapolate our results to the very low positive end of each trait distribution. Studies employing measures that are sensitive to the positive, low end of the distribution are needed^40,41^.

Specific limitations of the genetic analyses include the modest sample sizes and associated low power of several of the discovery GWAS analyses used to derive PRS; for certain phenotypes, the limited sample size may have led to less robust results (in particular for ANX and BD). Also, the estimates of variance explained by the PRS were modest; although these results indicate the presence of associations between genetic risk for disorders and traits, the degree to which common genetic risks are shared is unclear. Future studies utilizing larger GWAS datasets will be needed to estimate genetic correlations from molecular genetic data.

To conclude, our results from two different analytic approaches largely converge to suggest that the genetic factors that predispose to psychiatric disorders are also associated with continuous variation in milder, characteristic traits of these disorders. This provides genetic support for the hypothesis that many psychiatric disorders can be considered as extreme manifestations of continuous traits in the general population.

## Acknowledgements & Funding

Dr Martin was supported by the Wellcome Trust (Grant No: 106047). The CATSS study has support from the Swedish Research Council for Health, Working Life and Welfare and the Swedish Research Council.

We would like to acknowledge and thank the following consortia which provided GWAS summary statistics: Psychiatric Genomics Consortium (ASD, ADHD, Tourette’s syndrome, BD, OCD, MDD and Schizophrenia Working Groups), Lundbeck Foundation Initiative for Integrative Psychiatric Research (iPSYCH), iPSYCH-Broad Workgroup, Anxiety NeuroGenetics STudy (ANGST), International OCD Foundation Genetics Collaborative & OCD Collaborative Genetics Association Studies, EArly Genetics and Lifecourse Epidemiology Consortium (EAGLE), Social Science Genetic Association Consortium (SSGAC), and the Cognitive Genomics Consortium (COGENT).

The GWAS data for Tourette’s syndrome were supported by grants from the Judah Foundation, the Tourette Association of America, NIH Grants NS40024, NS016648, the American Recovery and Re-investment Act (ARRA) Grants NS040024-07S1, NS16648-29S1, NS040024-09S1, MH092289; MH092290; MH092291; MH092292; R01MH092293; MH092513; MH092516; MH092520; and the New Jersey Center for Tourette Syndrome and Associated Disorders (NJCTS).

The GWAS data for OCD were supported by grants from the Judah Foundation, the Tourette Association of America, NIH grants MH079489 and MH073250, American Recovery and Reinvestment Act (ARRA) awards NS40024-07S1, NS16648-29S1, MH071507, MH079489, MH079487, MH079488 and MH079494.

## Disclosures

Prof Larsson has served as a speaker for Eli-Lilly and Shire and has received research grants from Shire; all outside the submitted work. Prof Lichtenstein has served as a speaker for Medice.

